# Fecal Contributors to an Urbanized Estuary Identified by a Multi-Species qPCR Approach

**DOI:** 10.64898/2026.10.07.757507

**Authors:** Penny H Demetriades, Ania Brown, Jessica L. Jones, Sinéad Ní Chadhain, Ruth H. Carmichael, Brandi Kiel Reese

**Affiliations:** Stokes School of Marine and Environmental Sciences, University of South Alabama, Mobile, Alabama, USA; Dauphin Island Sea Lab, Dauphin Island, Alabama, USA; Gulf Coast Seafood Laboratory, U.S. Food and Drug Administration, Dauphin Island, Alabama, USA; Division of Seafood Safety, U.S. Food and Drug Administration, College Park, Maryland, USA; Department of Biology, University of Arkansas at Little Rock, Little Rock, Arkansas, USA

## Abstract

Coastal waters often experience fecal contamination from human (e.g., septic systems, wastewater treatment catchments and outfalls) and non-human (e.g., wildlife, domesticated pets, and livestock) origins, posing human health risks through contaminated fisheries and recreational waters. Quantitative PCR (qPCR) methods can identify and quantify fecal origins for their management by targeting *Bacteroides*, a dominant group of gastrointestinal bacteria with high host specificity. This study identified major contributors and drivers of fecal contamination to Mobile Bay, Alabama, an urbanized estuary in the southern United States. We utilized a multi-species qPCR source tracking approach targeting total *Bacteroides* and human-, pig-, dog-, cow-, and fowl-associated *Bacteroides*. Water samples and environmental parameters were collected over one year from 13 sites that represent a range of potential fecal inputs common to coastal waters. Human-associated feces were a consistent major contributor to the system (positive in 82% of samples), with localized inputs from pig (wild or domestic) and dog. Freshwater discharge and nitrogen constituents were good predictors of *Bacteroides* concentrations in this system, while areas with impervious surfaces and developed land cover may be contributing elevated fecal indicator levels. The multi-species qPCR source tracking approach described in this study can be implemented in other watersheds to support the development of better detection, identification, and fecal management strategies for community health of coastal areas.

**Importance:** (<150 words; nontechnical explanation of the significance of the study to the field) Humans, wildlife, domesticated animals, and livestock may contribute to fecal contamination of coastal waters. Fecal contamination can pose human health risks by making fisheries and recreational areas unsafe. To successfully mitigate fecal contamination, the origin (biological source) of feces needs to be known. We identified fecal origins to an urbanized estuary using a DNA-based approach that targets multiple potential fecal contributors through their associations with gut bacteria (*Bacteroides*). Our data set a baseline for fecal origins and concentrations in the Mobile Bay estuary and represent the most spatially and temporally comprehensive microbial source tracking effort in the region to date. Due to robust simultaneous cross-validation, this study also represents the first successful application of this source tracking approach for analysis of five species from the same sample. This approach is broadly transferrable to inform management and improve community and ecosystem health.

## Introduction

Coastal zones and shoreline systems are highly dynamic and support a disproportionately large fraction of the U.S. population relative to their small land cover (Reimann et al. 2023). Coastal areas receive and drain influxes of water from inland areas, and the urbanization of coastlines allows for the introduction of contaminants from various terrestrial origins into aquatic systems (MacDonnell et al. 2017). Fecal contamination is a recurring problem in coastal areas worldwide, especially those with developed land cover (Mallin et al. 2001, Tuholske et al. 2021, Fulke et al. 2025). Pathogens can be conveyed to waterways from both human (e.g., wastewater treatment collection systems and outfalls, septic systems) and non-human (e.g., wildlife, domesticated pets, livestock) origins (Fong & Lipp 2005, Li et al. 2022). For example, feces can be contributed through direct contact between wild or domesticated animals and waterways, improper land application of manure-based fertilizers, sewage and septic spills and leaks, and runoff from major rain events (Yan & Sadowsky 2006, Åström et al. 2013, Zhang et al. 2014). Globally, microbial inputs to coastal areas remain largely undefined (García Aljaro et al. 2018). A better understanding of fecal origin(s) is necessary to inform system-scale management that improves community and ecosystem health.

Waters receiving influxes of fecal contamination pose various health concerns for coastal residents, tourists, and shellfish consumers (Martinez-Manzanares et al. 1992, Pancorbo & Barnhart 1992, de Souza et al. 2019). Coastal areas that are under supported may have aging infrastructure that is prone to failure and higher frequencies of contamination events (Brown & Peterson 2002). Sewage spills are a common problem that can result in fishing area, beach, or swimming water closures. Closures are based on indicator organisms associated with fecal contamination in order to prevent illnesses from pathogens commonly found in feces, such as *Vibrio*, *Salmonella*, *Shigella*, and *Campylobacter* (Rose et al. 2001, Savichtcheva & Okabe 2006, McKee & Cruz et al. 2021). Gastrointestinal illnesses may be a result of ongoing and emerging infectious diseases from human and non-human fecal contributors that are largely undefined (García-Aljaro et al. 2018, Ramsden et al. 2023). It is important to distinguish between human and non-human fecal contributions to mitigate contamination sources for the protection of public health. Human fecal contributions may pose a distinct risk from other animal feces due to differential occurrences of pathogens. Waterborne pathogens and the illnesses they cause comprise a major worldwide public health concern (García-Aljaro et al. 2018, ASM 2025, WHO 2026). Water quality degradation, therefore, poses a significant threat to human health, coastal resources, and coastal economies.

Microbial source tracking (MST) is a technique for detecting fecal contamination in the environment and identifying its origin(s). The methods used for MST have evolved since the start of the field, and methods differ in capability and rigor. Traditional MST methods target fecal indicator bacteria (e.g., *E. coli*, *Enterococcus*) abundance and phenotypes through plate-based, classical microbiology methods (Raza et al. 2021). These traditional methods can detect the presence of fecal-associated bacteria in the environment, but they are not well-suited to determine the absolute abundance of bacteria in the environment or their biological sources (Scott et al. 2002, Simpson et al. 2002, Raza et al. 2021). The inability of traditional microbial source tracking methods to distinguish between biological sources leads to difficulty in managing areas that exhibit elevated levels of fecal indicator bacteria during water quality monitoring. Additionally, recent research has suggested that the organisms targeted by classical MST methods can adapt and persist in the natural environment, becoming “naturalized,” and therefore decreasing their reliability as indicators of fecal contamination (Ishii & Sadowsky 2008, Ferguson & Signoretto 2011).

Quantitative PCR (qPCR) was first introduced as an MST tool in the 2000s, and its use has increased with the development of new target assays (Reischer et al. 2007, Raza et al. 2021). Host-associated *Bacteroides* are commonly used targets for source-specific qPCR. *Bacteroides* are anaerobes present in high concentrations in feces, and they can be successfully used to distinguish between hosts due to their high host specificity (Layton et al. 2006). Host-associated *Bacteroides* have exhibited rapid decay rates outside of their host environment (Dick et al. 2010) and may, therefore, serve as a more reliable indicator of recent fecal contamination compared to traditional indicators (Cloutier & McLellan 2017). The ability of qPCR technology to identify the origin(s) of fecal contamination and provide a quantitative measure of target organism abundance can allow for origin-specific management of contaminated areas. However, the utility of qPCR as a source tracking tool is restricted by issues of cross-reactivity, sensitivity, and specificity. The efficacy of genetic markers for MST is limited by factors such as geographic location, host diet, and climate (Ahmed et al. 2016, Raya et al. 2024). Primer sets should, therefore, be assessed on a watershed-by-watershed basis to evaluate their performance and confirm a lack of cross-reactivity with other fecal markers.

Similar to many urbanized coastal areas worldwide, the Mobile Bay estuary and its tributaries have a history of recurring fecal contamination. This is reflected in the health of the coastal community. In recent years, human and wildlife-associated transmissible pathogens have caused hundreds of cases of illnesses in Alabama residents (ADPH 2021). Areas that represent 80% of the current total oyster farming capacity in the state and nearly $2 million in local revenue have been subject to closures due to high levels of microbial contamination from undefined origins (Grice et al. 2017). Currently, a majority of Alabama’s waterways are included on the Alabama Department of Environmental Management (ADEM) 303(d) list of pathogen-impaired waters.

To date, MST methods have not been widely applied in the southern United States. Previous source tracking studies in Alabama have consisted of small-scale spatially and temporally discrete projects with limited application for system-scale management. These studies have also been limited in their methodologies, specifically with respect to their lack of range in targets (*E. coli*, *Enterococci*, general *Bacteroides*, human-associated *Bacteroides*, and cow-associated *Bacteroides*) for fecal origin identification (Ernest et al. 2011, Xue et al. 2018, Frith et al. 2021). Furthermore, many fecal pathogens, including those of potential wildlife origin, are not routinely tested for in Alabama. Therefore, the species-specific contribution of local wildlife to microbial indicators in Alabama waters remains largely unknown (Pandey et al. 2014). Verification of a set of genetic markers that target a range of hosts has not yet been established for the Mobile Bay estuary and is pertinent for proper management of fecal contamination for human health and safety.

The objective of this study was to evaluate the applicability of multi-species qPCR methods to inform reliable MST efforts that are transferable to urbanized estuaries globally where fecal contamination is a concern. These data set a baseline for fecal origins and concentrations in an urbanized estuary and represent the most spatially and temporally comprehensive MST effort in the region to date. To our knowledge, this study is the first conventional qPCR source tracking effort to simultaneously validate five species-specific markers for use on the same sample. This work provides vital new data to better understand the origins of fecal contamination to coastal areas and inform management that improves community and ecosystem health.

## Materials and Methods

### Study Sites

Mobile Bay is a shallow urbanized estuary on the southern coast of the United States with the fourth largest freshwater discharge among continental U.S. watersheds (Du et al. 2018). The mean discharge for Mobile Bay (1,677 m3/s) typically exhibits seasonality through higher freshwater discharge in winter and spring and lower discharge in summer and fall (Stumpf et al. 1993, MBNEP 2012, Du et al. 2018).

Sample collection occurred at 13 sites surrounding Mobile Bay (**Fig. 1** and **Table 1**) that represent a range of potential microbial inputs to coastal Alabama (including human, wildlife, domesticated pets, and livestock). West Mobile Bay (WMB) samples were collected at the outfall pipe of the Mobile Area Water and Sewer System (MAWSS) wastewater treatment plant and represent undiluted treated effluent.

**Figure 1.**
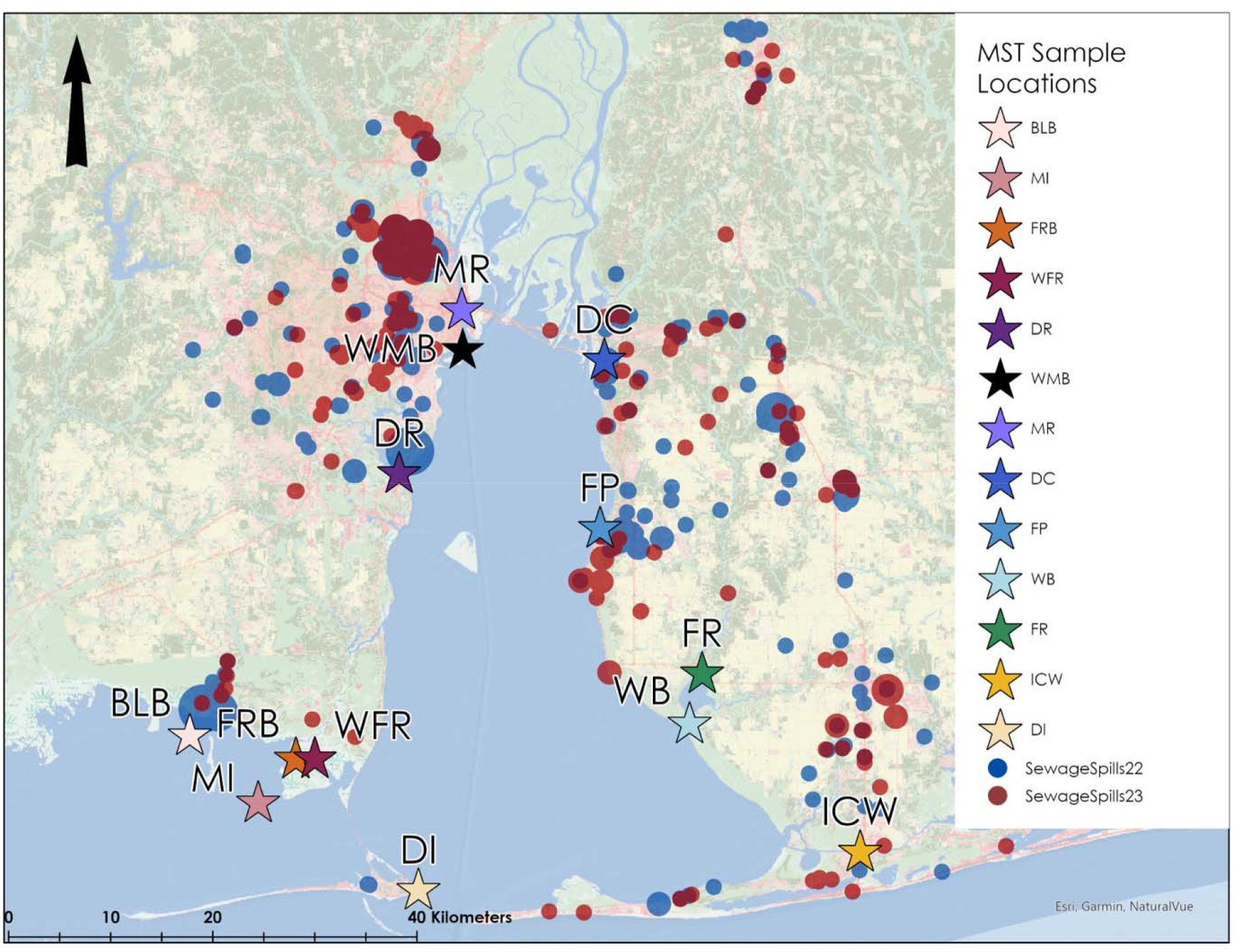
Location of the 13 sampling sites and known ADEM sewage spills in the area (2022-2023). Basemap: USGS National Land Cover Database (NLCD).

**Table 1.** Latitude and longitude (decimal degrees), land use designation, percent urban development of basin, and average water depth, temperature, and salinity of the 13 sampling sites. Average water depth, temperature, and salinity were collected a handheld YSI during sampling. Land use designations were determined using the USGS National Land Cover Dataset and basin percentage of urban development and impervious cover were calculated using USGS StreamStats.

| Site | Name | Latitude | Longitude | Land Use<br>Designation | % of Basin<br>with Urban<br>Developme<br>nt | % of<br>Basin with<br>Imperviou<br>s Cover | Average<br>water<br>depth<br>(m) | Average<br>tempera<br>ture<br>(°C) | Average<br>salinity |
| --- | --- | --- | --- | --- | --- | --- | --- | --- | --- |
| 1<br>(BLB<br>) | Bayou<br>la Batre | 30.36783 | -88.27678 | Open water | 11.8 | 2.9 | 2.3 | 24.2 | 17.5 |
| 2<br>(MI) | Marsh<br>Island | 30.32111 | -88.21825 | Open water | 0 | 0 | 1.1 | 25.2 | 15.9 |
| 3<br>(FRB) | Fowl<br>River<br>Bay | 30.35499 | -88.19289 | Open water | 4.6 | 1.3 | 1.3 | 24.4 | 15.3 |
| 4<br>(WFR<br>) | West<br>Fowl<br>River | 30.36841 | -88.16145 | Emergent<br>herbaceous<br>wetland | 6.43 | 2.1 | 1.9 | 24.5 | 11.7 |
| 5<br>(DR) | Dog<br>River | 30.56683 | -88.09254 | Developed,<br>medium<br>intensity | 53.4 | 16.2 | 3.0 | 24.1 | 5.0 |
| 6<br>(WM<br>B) | West<br>Mobile<br>Bay<br>(WTP<br>effluent<br>) | 30.66044 | -88.03650 | Developed,<br>medium<br>intensity | 74.3 | 55.0 | 1.7 | 25.9 | 0.5 |
| 7<br>(MR) | Mobile<br>River | 30.69085 | -88.03730 | Developed,<br>high<br>intensity | 89.8 | 76.7 | 5.6 | 23.4 | 6.5 |
| 8<br>(DC) | D'Oliv<br>e Creek | 30.65269 | -87.91223 | Developed,<br>medium<br>intensity | 39.4 | 11.2 | 2.9 | 23.2 | 0.5 |
| 9 (FP) | Fairhop<br>e Pier | 30.52499 | -87.91552 | Developed,<br>medium<br>intensity | 61.4 | 19.2 | 2.3 | 23.8 | 8.1 |
| 10<br>(WB) | Weeks<br>Bay | 30.37663 | -87.83694 | Developed,<br>low<br>intensity | 10.3 | 2.7 | 2.3 | 24.0 | 8.9 |
| 11<br>(FR) | Fish<br>River | 30.41475 | -87.82583 | Emergent<br>herbaceous<br>wetland | 10.6 | 2.9 | 1.8 | 24.2 | 4.9 |
| 12<br>(ICW<br>) | Intraco<br>astal<br>Waterw<br>ay | 30.27995 | -87.68583 | Developed,<br>medium<br>intensity | 17.6 | 5.2 | 3.3 | 24.5 | 14.2 |
| 13<br>(DI) | Dauphi<br>n Island | 30.25016 | -88.07558 | Developed,<br>low<br>intensity | - | - | 0.7 | 22.9 | 15.6 |

### Sample Collection

From September 2022 to September 2023, water samples were collected from all 13 sites every four weeks. Sample collection occurred between high tide and low tide to capture microbial contamination as it flushed out of the bay. Water samples for qPCR analyses were collected in sterilized 10L carboys filled just below the surface of the water and placed immediately on ice for transport. Water samples for nutrient analysis, stable isotopes, and trace metals were sampled using a horizontal water sampler at 0.5m below the water surface. Nutrient, stable isotope, and trace metal samples were pre-filtered through a 200μm mesh screen and collected in acid-washed 500mL bottles.

Physicochemical parameters (i.e., salinity, pH, water temperature, and dissolved oxygen) of the water were measured at the location of sample collection with a handheld YSI Pro 2030 during each sampling event. Wind speed and air temperature were collected at each site with a Kestrel 3000 Weather Meter. Water depth at each site was measured with a portable depth sounder (Speedtech). Data for rainfall, freshwater discharge, and flow rates were compiled from community partners and public databases (South Alabama Mesonet, United States Geological Survey (USGS)). Land use designations were determined using the USGS National Land Cover Dataset. Basin percentage of urban development and impervious cover were calculated using USGS StreamStats. Two-week average turbidity and wind speed measurements were taken from Alabama’s Real-Time Coastal Observing System (ARCOS), which collects turbidity data every 30 minutes and wind speed every minute.

### Sample Processing

Water samples for nutrient analysis, stable isotopes, and trace metals (including Sr:Ba as a salinity proxy) were analyzed via established methods as described in Brown 2025. Water samples for molecular analysis were filtered immediately upon returning to the lab (within 6 hours of collection). Water was filtered until refusal through 47mm 0.22μM Millipore Nitrocellulose filters (Burlington, Massachusetts) in triplicate for qPCR analysis. In-line filter towers for the filters were connected to a peristaltic pump via sterilized tygon tubing. Between sites, tubing was switched out and filter towers were sterilized with 70% ethanol. The volume that passed through each filter was measured and recorded for use in qPCR quantification of targets. All filters were folded with biomass facing biomass and stored at -80^°^C until extraction. Total DNA was extracted from each of the triplicate water filters with the Qiagen PowerWater DNA Extraction Kit (Venlo, Netherlands), following the manufacturer’s instructions. Each extraction used ½ of the 47mm filter. All extractions were conducted in a biosafety cabinet with PPE (face mask, bouffant cap, shoe covers, and lab coat), instruments were flame sterilized between samples, and blanks were run with each set of extractions. Extracted DNA was quantified with the Qubit Broad Range (BR) DNA quantification kit (Invitrogen, Waltham, Massachusetts).

### Quantitative PCR

Multiple samples of fresh chicken, duck (wild and domestic), dog, and cow fecal samples from the Mobile Bay area were collected in sterilized containers to serve as positive controls for the chicken/duck, dog, and cow targets. Wastewater influent from the MAWSS wastewater treatment plant in Mobile, AL was used as the human positive control. Six feral hog intestines (collected with the help of Grand Bay National Estuarine Research Reserve) were used as the pig positive control. Positive control DNA was extracted from fecal samples using the Zymo Quick-DNA Fecal/Soil Microbe Kit (Irvine, California). Primer sets (**Table 2**) were verified against positive controls from fecal samples from all host targets to ensure there was no cross reactivity between markers in the Mobile Bay watershed. Fecal samples for the positive controls were amplified with their respective primer sets and cloned into *E. coli* using the Invitrogen TOPO TA Cloning Kit. Clones were Sanger sequenced at Eurofins USA and checked for correct target amplification with NCBI BLAST. To generate sequences for gene blocks (used as qPCR standards for quantification), short sequences of the T7 promoter were added to the 5’ and 3’ ends of the positive controls (inclusive of forward primer and reverse primer) to buffer against end degradation. Gene blocks were ordered from Integrated DNA Technologies (IDT), Coralville, Iowa. Quantitative PCR (qPCR) targeted *Bacteroides* associated with human (HF183/BacR287), dog (DG3), pig (Pig-2-Bac), fowl (Chicken/Duck-Bac), and cow (CowM2), and general *Bacteroides* (AllBac) in all water samples. qPCR assay master mixes (**Table A1**) and cycling conditions (**Table A2**) were optimized for each primer set and incorporated an internal amplification control (IAC). The IAC was used to assess amplification inhibition. All reactions were run on Applied Biosystems 7500 Real-Time PCR Instruments (Waltham, Massachusetts). Replicate extractions for each sample were pooled before being run in triplicate (technical replicates) for each qPCR assay. Triplicate no template controls (NTC) and triplicate standard curves were included in all runs. A manual threshold setting for each assay was determined based on amplification plots.

**Table 2.**
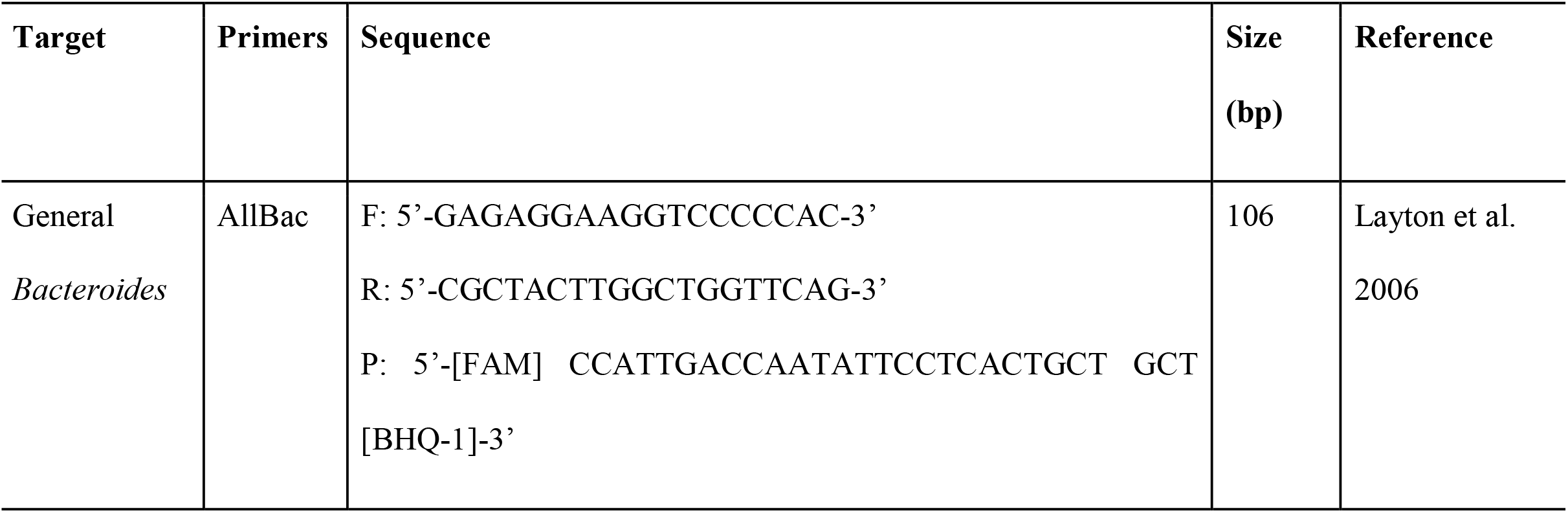
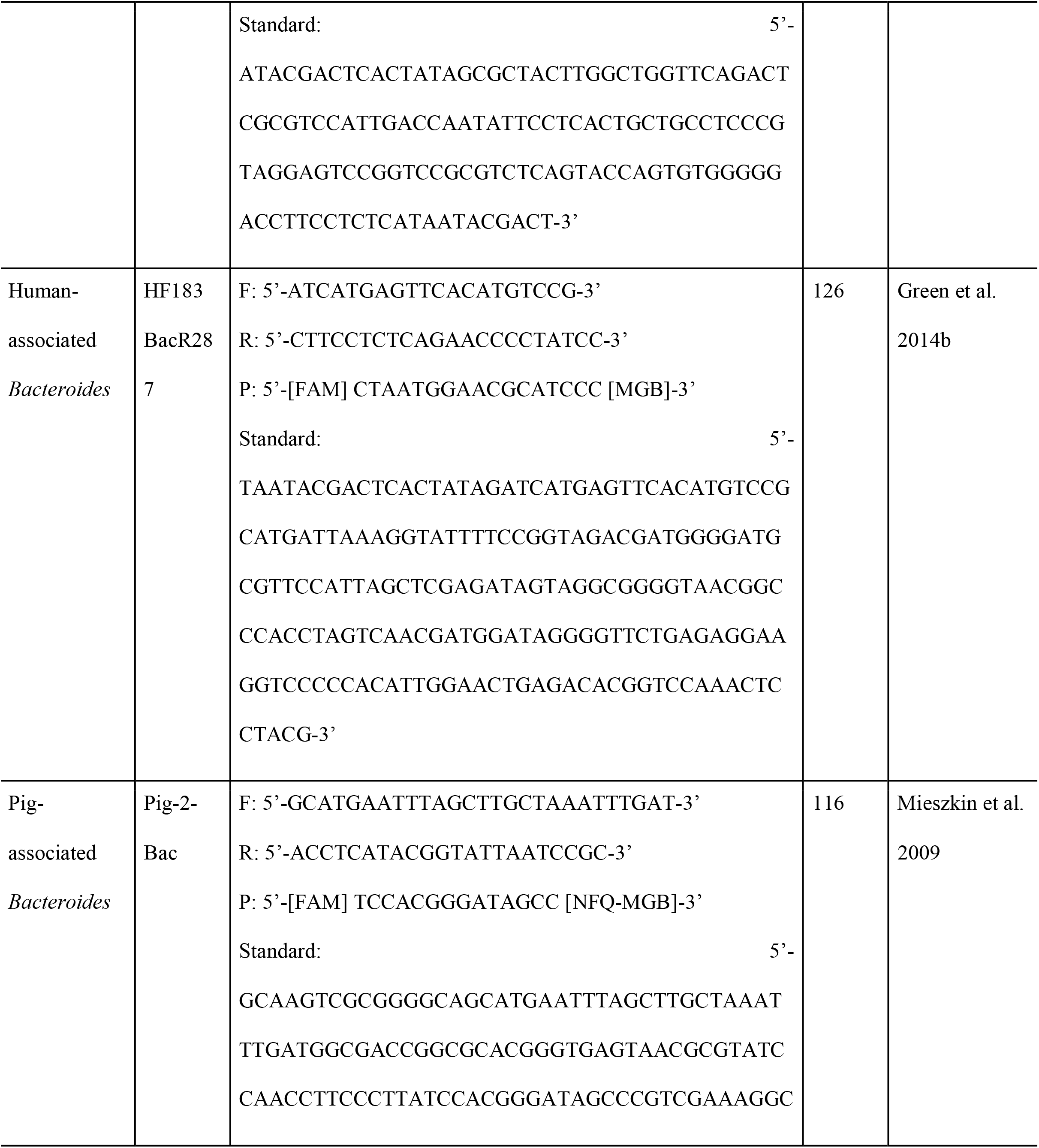
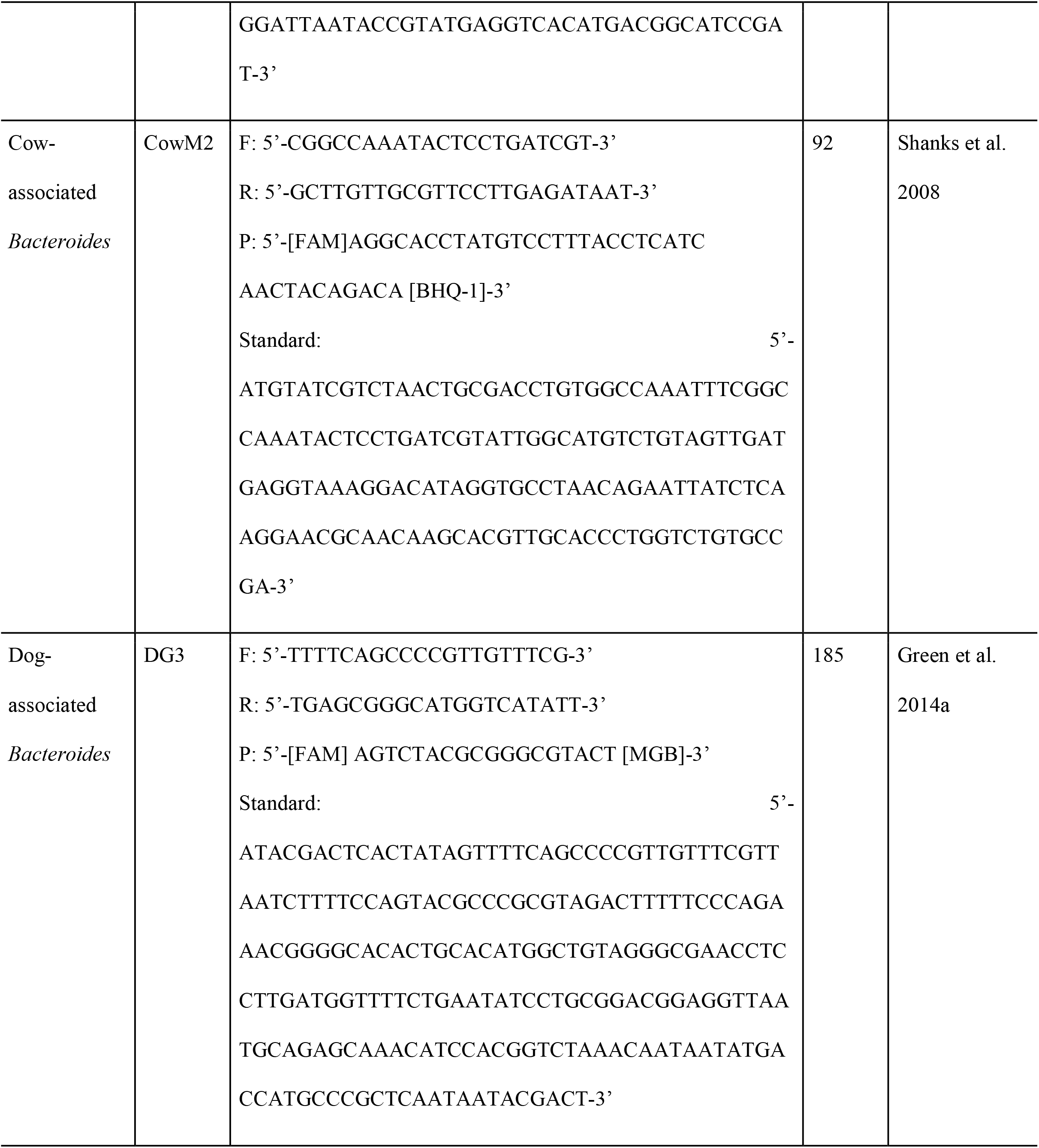
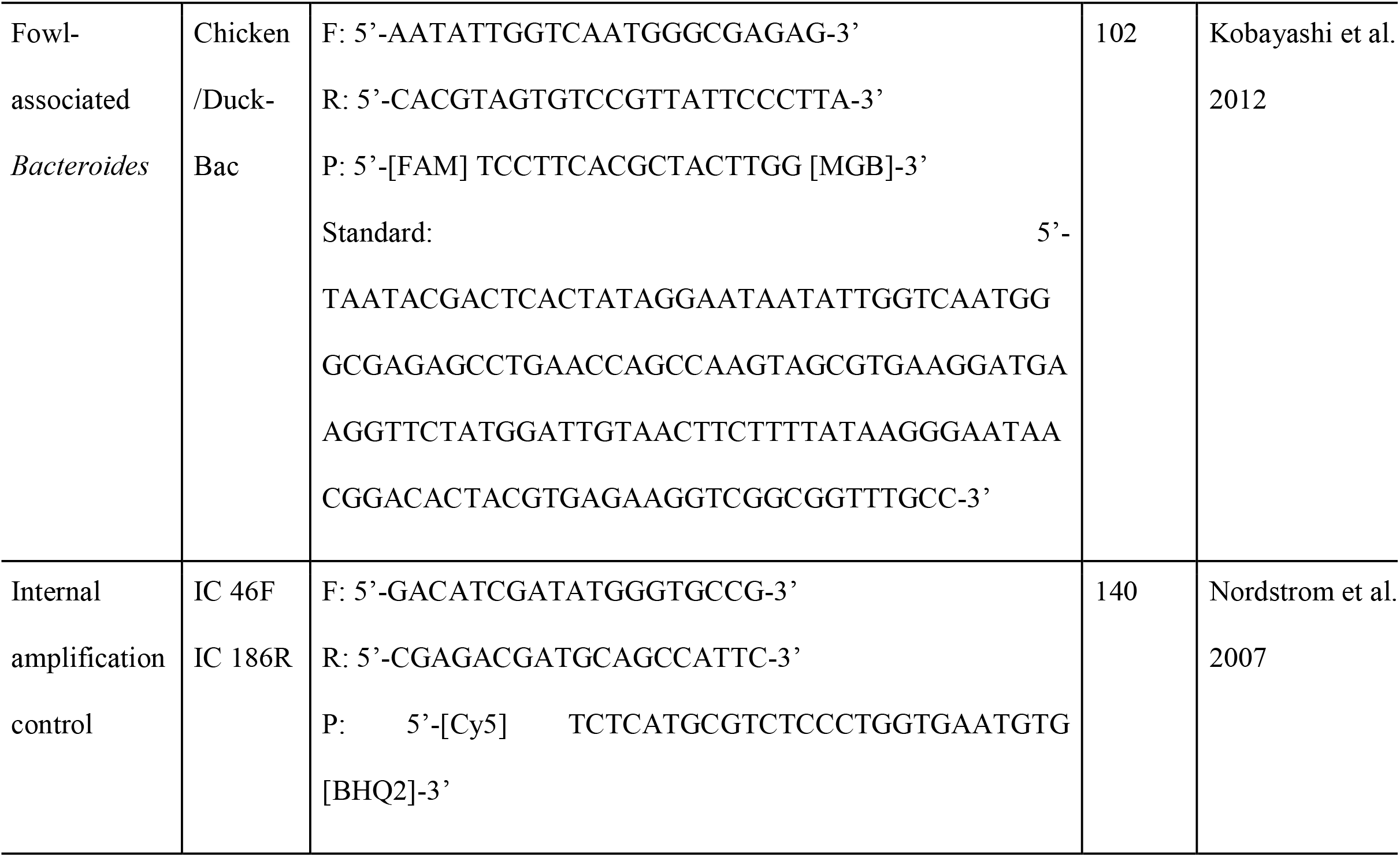
Primer, probe, and standard sequences for qPCR assays. Forward primer listed as “F,” reverse primer listed as “R,” and probe listed as “P.” Sequences from the fecal positive control PCR products were used to create gene block standards (Integrated DNA Technologies, California).

### Data Analysis

Amplification curves were manually screened to confirm instrument determinations before quantified outputs were used to determine copy numbers. Quantities of each *Bacteroides* target were determined using qPCR standard curves and then converted to copies per 100mL based on the volume of water processed for each filter. Technical replicates were averaged to generate a single quantity per sample for each qPCR marker. The limit of detection (LOD) was determined for each assay based on the lowest detected quantity for all three replicates in the standard curve. To avoid zero-inflation during statistical analyses, non-detect samples were calculated as half of the LOD based on the volume of water processed for each sample. No potential outlying qPCR replicates were removed.

All qPCR and environmental data were compiled into Excel. Statistical analyses were conducted using R version 4.3.2 (R Core Team) in RStudio, and plots were generated with ggplot2. Marker-to-marker concentrations were compared with Student’s t-tests, and correlation analysis (Pearson’s) was used to determine relationships between the species-specific and AllBac markers. A stratified statistical approach (separated by qPCR marker) using analysis of variance (ANOVA) regression was used to indicate which environmental parameters were significant (p<0.05) drivers of *Bacteroides* marker copy numbers. The Multi-Model Inference (MuMIn) model.sel function was used to indicate which environmental parameter was the best predictor of average *Bacteroides* copy number and to determine the best temporal model (ARCOS environmental parameters averaged over two weeks, one week, two days, or one day) for the relationships between turbidity or windspeed and average copy number. A dredge function model selection was also used to determine the best environmental predictor of *Bacteroides* concentrations. The vegan community ecology package was used to construct non-metric multidimensional scaling (NMDS) ordinations and Principal Components Analysis (PCA). The NMDS was generated with Bray-Curtis. NMDS and PCA were visualized with plots.

## Results

A total of 180 water samples were analyzed in triplicate for each of the six qPCR assays (all *Bacteroides*, human-, pig-, dog-, cow-, and fowl-associated *Bacteroides*). All qPCR runs had an efficiency between 90% and 110% and r^2^ > 0.99. LOD for the assays were between 1.7x10^1^ and 5.2x10^1^ gene copies per reaction. NTC and extraction blanks had no amplification. Internal amplification controls did not show amplification inhibition.

Extraction efficiency of the PowerWater kit was determined to have a consistent average recovery (ranging from 4.6% to 12.7%) across ten different filter halves from the same water sample. Therefore, variations in samples are assumed to not be significantly affected by differential extraction efficiencies.

### General Trends

The general *Bacteroides* marker (AllBac) was consistently detected in the samples at higher levels than the other (species-specific) *Bacteroides* markers (p=0.01, **Fig. 2**). The AllBac marker had the highest number of positive samples, followed by human and pig, with all other markers having <50% positive samples (**Table 3**). Correlation analysis showed a positive relationship between human-associated *Bacteroides* and AllBac (p<0.001, r^2^=0.696) (**Fig. A1**). The fowl marker also had a moderately positive correlation with AllBac (p<0.001, r^2^=0.597) (**Fig. A2**). Although the AllBac marker was included to serve as a measure of total fecal contamination, the combined species-specific (human, pig, dog, fowl, cow) marker copy numbers on average accounted for 6.9 ± 1.0% of the AllBac gene copy numbers, with one exception of a species-specific marker accounting for >100% of the AllBac copy numbers at WMB in July 2023.

**Figure 2.**
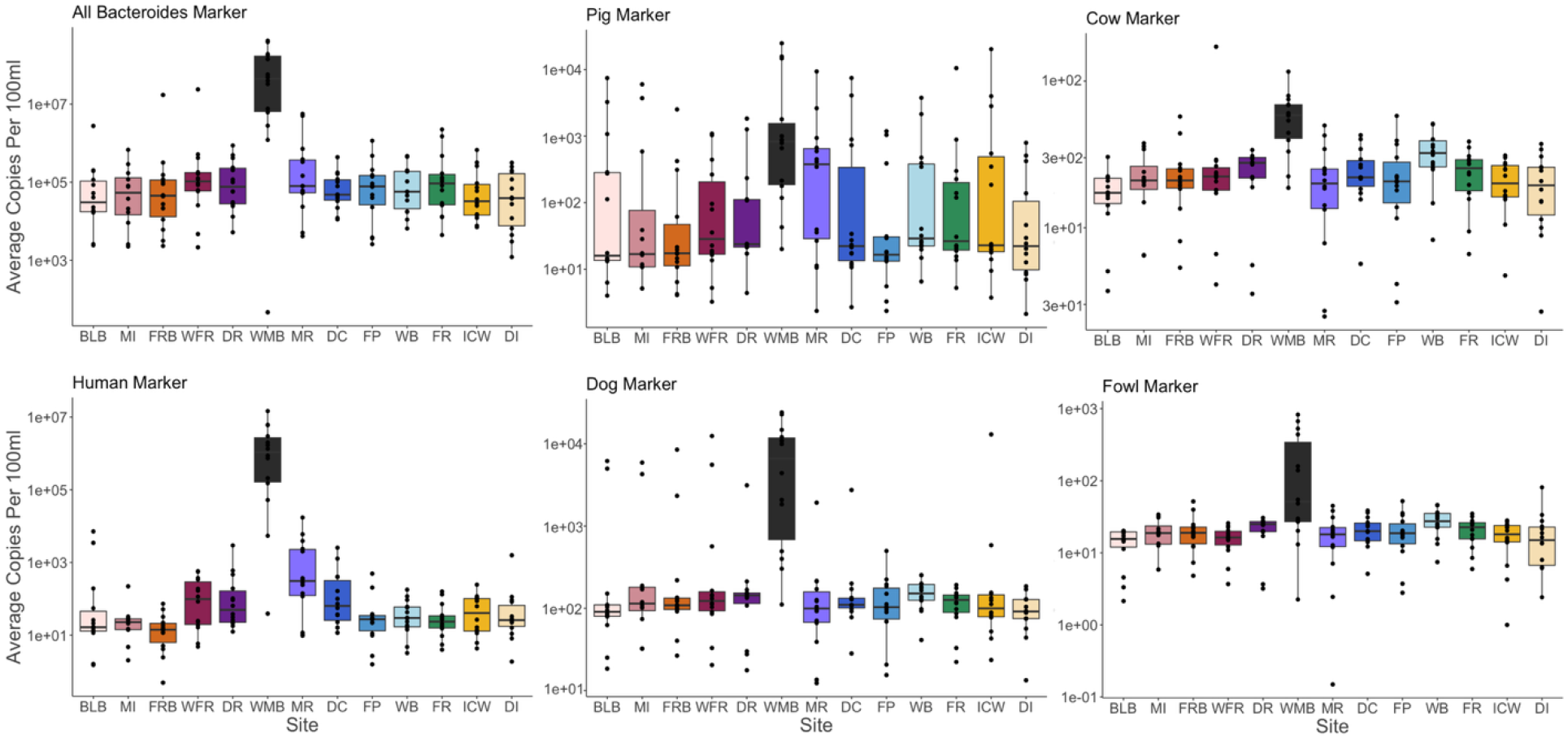
Average *Bacteroides* gene copy numbers for each marker over all time points. Note differences in Y-axis scales across panels.

**Table 3.** Percent of samples where marker was detected (positive) and mean concentration of each marker across samples where detection was positive (in gene copies (gc)/100mL).

|  | AllBac | Human | Pig | Dog | Fowl | Cow |
| --- | --- | --- | --- | --- | --- | --- |
| Positive samples | 99% | 82% | 58% | 24% | 20% | 1.1% |
| Mean (gc/100mL) | $8.9 \times 10^6$ | $2.05 \times 10^5$ | $9.72 \times 10^2$ | $7.33 \times 10^4$ | $1.65 \times 10^1$ | $1.03 \times 10^1$ |

**Table 4.** Relationships (ANOVA regression analysis, p<0.05) between environmental parameters and *Bacteroides* gene copy numbers for all sites (WMB included). Positive relationships are denoted with a (+) and negative relationships are denoted with a (-). No symbol indicates no statistical relationship.

|  | AllBac | Human | Pig | Dog | Fowl | Cow |
| --- | --- | --- | --- | --- | --- | --- |
| Salinity | - | - |  |  | - | - |
| TDN | + | + | + | + | + | + |
| $\text{NH}_4^+$ | | | | | + | + |
| $\text{NO}_3^-$ | + | + | + | + | | + |
| $\text{NO}_2^-$ | | | | | + | |
| Water Temp | + |  |  |  |  |  |
| pH | - |  |  | + | + | + |
| DO | - |  |  | + |  |  |
| Rainfall |  |  |  |  | + | + |
| Turbidity | + | + |  |  |  |  |
| Discharge |  | + |  |  |  |  |
| Urban Development | + | + | + | + |  |  |
| Impervious Cover | + | + | + | + |  |  |
| $\delta^{15}\text{N}$ | - | - | - | | - | - |
| Sr:Ba |  | - |  |  | - | - |

Average *Bacteroides* copy numbers for all markers differed between sites, despite intra-site variation. There were higher overall *Bacteroides* concentrations at the WMB site (wastewater effluent) than any other site (p<0.001). Temporally, human inputs were more consistent compared to inputs from dog and pig, which exhibited elevated marker concentrations at some timepoints (**Fig. 2**). There was a positive correlation between human-associated *Bacteroides* and AllBac (total) and the two markers displayed similar variation among sites and sampling dates. However, some sites were, on average, dominated by other species-specific origins (**Fig. 3**). Dog-associated *Bacteroides* were prominent in the Eastern Mississippi Sound sites (MI, FRB, and WFR), and pig-associated *Bacteroides* contributions were prominent on the eastern coast of Mobile Bay (DC, FP, WB, FR, ICW). There were no seasonal trends in *Bacteroides* across sites (**Fig. A3**).

**Figure 3.**
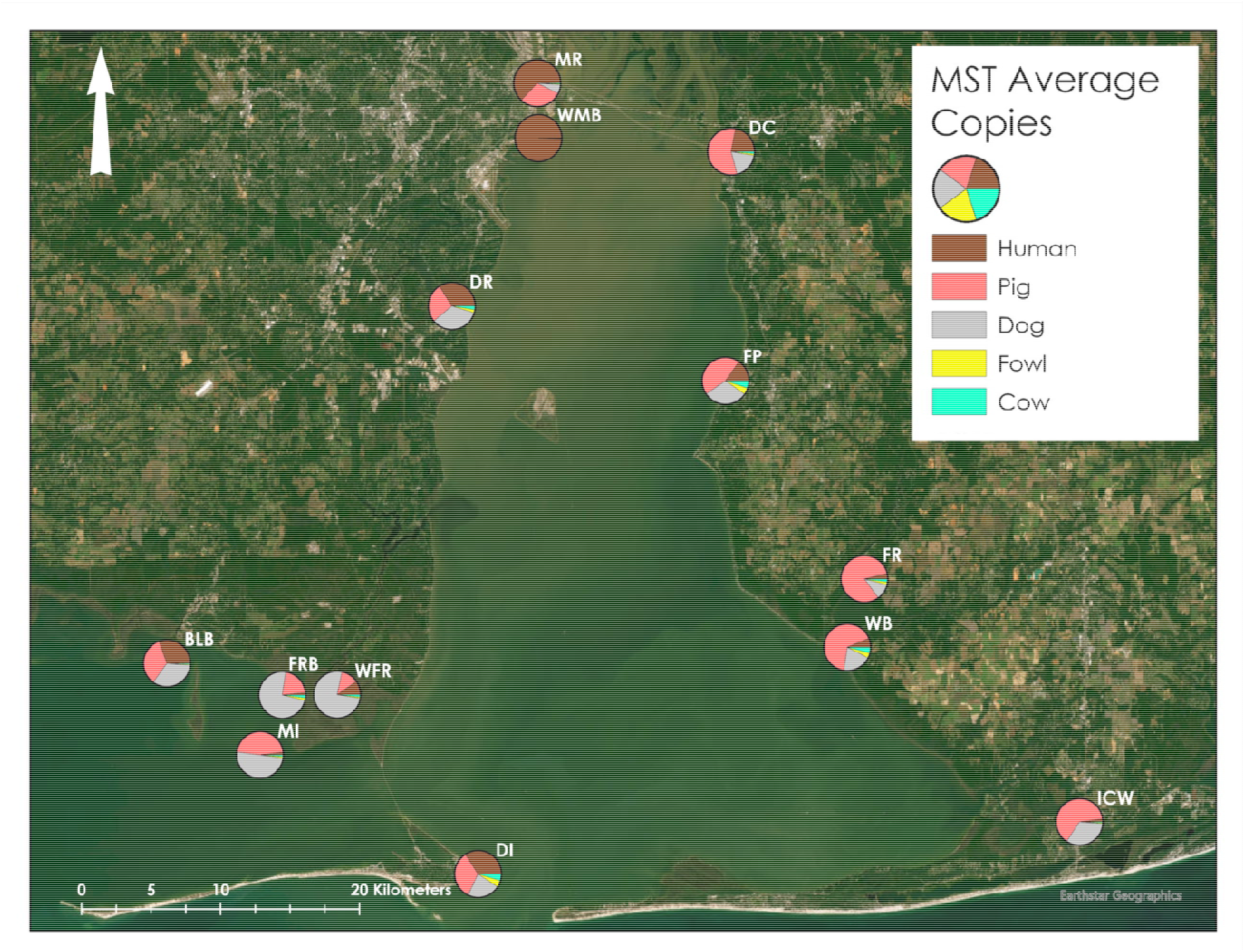
Average gene copies of species specific *Bacteroides* markers (human, cow, dog, pig, and fowl) at each site over all timepoints. On average, the combined species-specific copy numbers accounted for 6.9% of the AllBac copy number.

### Marker-specific Trends

#### All *Bacteroides* (AllBac)

Regression analysis showed positive relationships between the general *Bacteroides* marker (AllBac) and water temperature (p<0.001), turbidity (p=0.002), total dissolved nitrogen (TDN, p<0.001), nitrate (NO ^-^, p=0.02), percent urban development (p<0.001), and percent impervious cover (p=0.01). There were negative relationships with salinity (p<0.001), Sr:Ba (p<0.001), dissolved oxygen (p<0.001), pH (p<0.001), and δ^15^N (p<0.001). For the AllBac marker, the WMB site (wastewater effluent) had higher copy numbers than any other site (p=0.001). When the WMB site is treated as an outlier and excluded from the statistical analysis, the relationships do not change, with the exception of land use designation having a relationship (p=0.001) and nitrate and δ^15^N no longer having a relationship with gene copy number.

#### Human

Regression analysis showed positive relationships between the human *Bacteroides* marker and turbidity (p=0.01), freshwater discharge (p=0.004), TDN (p<0.001), nitrate (p<0.001), percent urban development (p<0.001), and percent impervious cover (p<0.001). There were negative relationships between human and salinity (p=0.04), Sr:Ba (p=0.01), and δ^15^N (p<0.001). There was a relationship (p<0.001) between land use designation and marker copy numbers, with medium intensity developed sites presenting higher average concentrations than other land use designations. The WMB site (wastewater effluent) had higher copy numbers than any other site (p<0.001). When the WMB site is treated as an outlier and removed from the statistical analysis, the relationships do not change, with the exception of a positive relationship with nitrite (p<0.001), instead of nitrate. Additionally, without WMB, there are no longer relationships with salinity, Sr:Ba, and δ^15^N.

#### Pig

Regression analysis showed positive relationships between the pig *Bacteroides* marker and TDN (p<0.001), nitrate (0.005), percent urban development (p=0.01), and percent impervious cover (p=0.01). There was a negative relationship with δ^15^N (p<0.001). There was a relationship between land use designation and marker copy numbers (p=0.04), with medium intensity developed sites presenting higher average concentrations than other land use designations. The ICW site had higher copy numbers than any other site (p<0.001). When the WMB site (wastewater effluent) is treated as an outlier and removed from the statistical analysis, the relationships do not change, with the exception of a relationship with pH (p=0.001) and no relationship with nitrate.

#### Dog

Regression analysis showed positive relationships between the dog *Bacteroides* marker and pH (p<0.001), dissolved oxygen (p=0.002), TDN (p<0.001), nitrate (p=0.02), percent urban development (p=0.04), and percent impervious cover (p=0.03). There were negative relationships with water temperature (p<0.001) and δ^15^N (p<0.001). The WMB site (wastewater effluent) had higher copy numbers than any other site (p<0.001). When the WMB site is treated as an outlier and removed from the statistical analysis, the relationships do not change, with the exception of new relationships with land use designation (p=0.03), ammonium (p=0.02), and nitrite (p=0.005). There are no longer relationships with TDN and δ^15^N.

#### Fowl

Regression analysis showed positive relationships between the fowl *Bacteroides* marker and pH (p<0.001), TDN (p<0.001), ammonium (p<0.001), and nitrite (p<0.001). There were negative relationships with salinity (p<0.001), Sr:Ba (p<0.001), and δ^15^N (p<0.001). The WMB site (wastewater effluent) had higher copy numbers than any other site (p<0.001). When the WMB site is treated as an outlier and removed from the statistical analysis, the relationships do not change, with the exception of a relationship with land use designation (p=0.003) and there no longer being any relationships with ammonium, TDN, or δ^15^N.

#### Cow

Regression analysis showed positive relationships between the cow *Bacteroides* marker and pH (p<0.001), TDN (p<0.001), ammonium (p=0.04), nitrate (p=0.003), and 24-hour average rainfall (p=0.007). There were negative relationships with salinity (p<0.001), Sr:Ba (p<0.001), and δ^15^N (p<0.001). When the WMB site (wastewater effluent) is treated as an outlier and removed from the statistical analysis, the relationships do not change, with the exception of new relationships with land use designation (p=0.002) and nitrite (p<0.001) instead of nitrate and TDN.

#### *Bacteroides* environmental parameter relationships

Model selection (MuMIn) analysis found TDN (AICc, weight=0.988, sum of squares (sum sq)=1022) was the most important environmental factor for predicting *Bacteroides* gene copy numbers. A different model selection (dredge) analysis found salinity was the most important factor (AICc, weight=0.97, sum sq=122) (**Table A2**). Two-week averages for the ARCOS data (turbidity weight=1; wind speed weight=0.387) were best correlated with *Bacteroides* gene copy number; compared to one-week, two-day, and one-day averages.

The NMDS ordination plots displayed strong site grouping (**Fig. 4**), with MR and FP (mid to north Mobile Bay) sites clustering together; FRB and WFR (adjacent Eastern Mississippi Sound sites) clustering; and WMB, DC, BLB, DI, and MI (wastewater outfalls and southern sites) clustering. The NMDS plot also showed strong clustering of *Bacteroides* gene copy numbers based on amount of freshwater discharge (**Fig. 5**).

**Figure 4.**
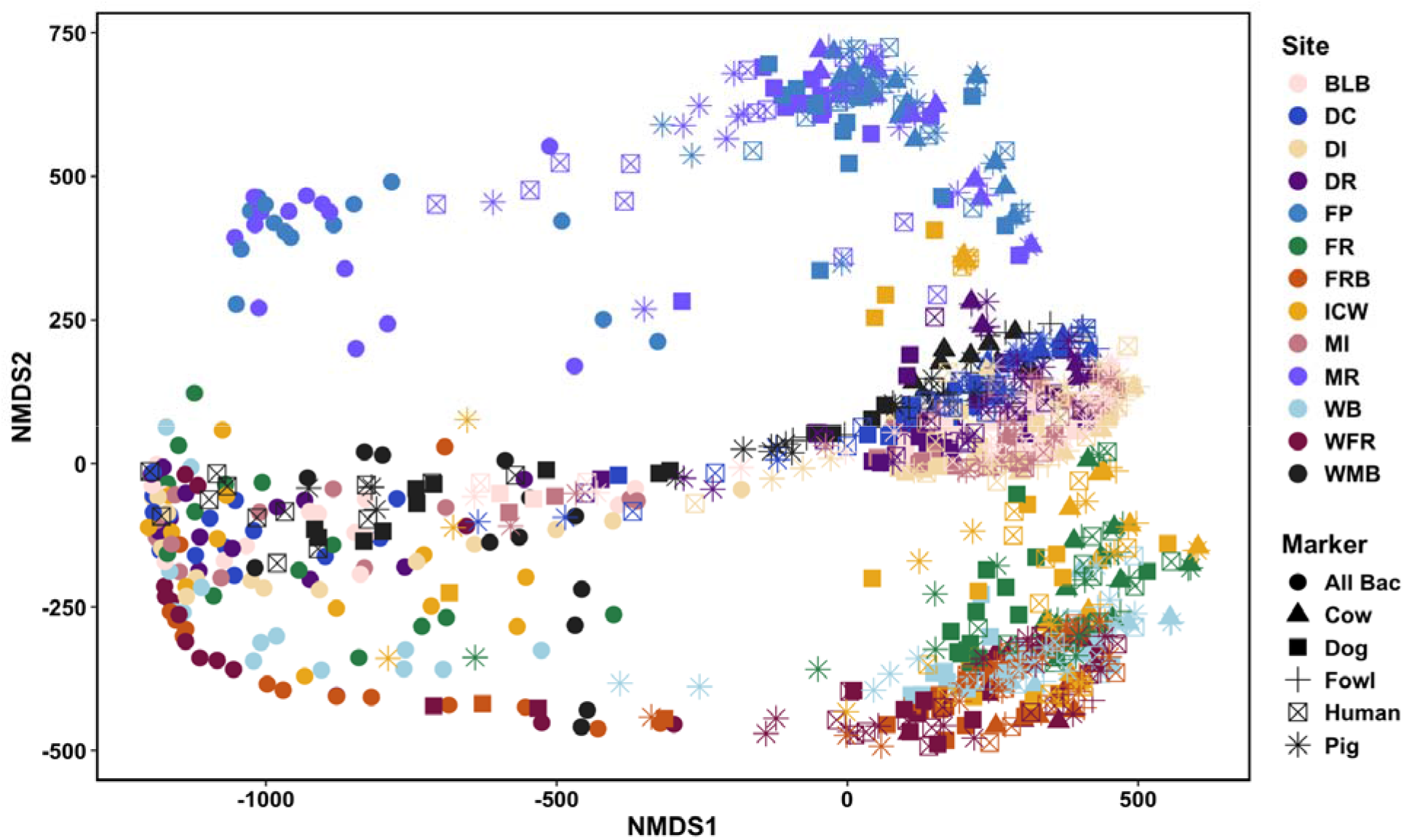
*Bacteroides* gene copy number ordinations (NMDS) constructed from environmental parameters and nitrogen measurements. Points are colored by site and shape denotes *Bacteroides* marker.

**Figure 5.**
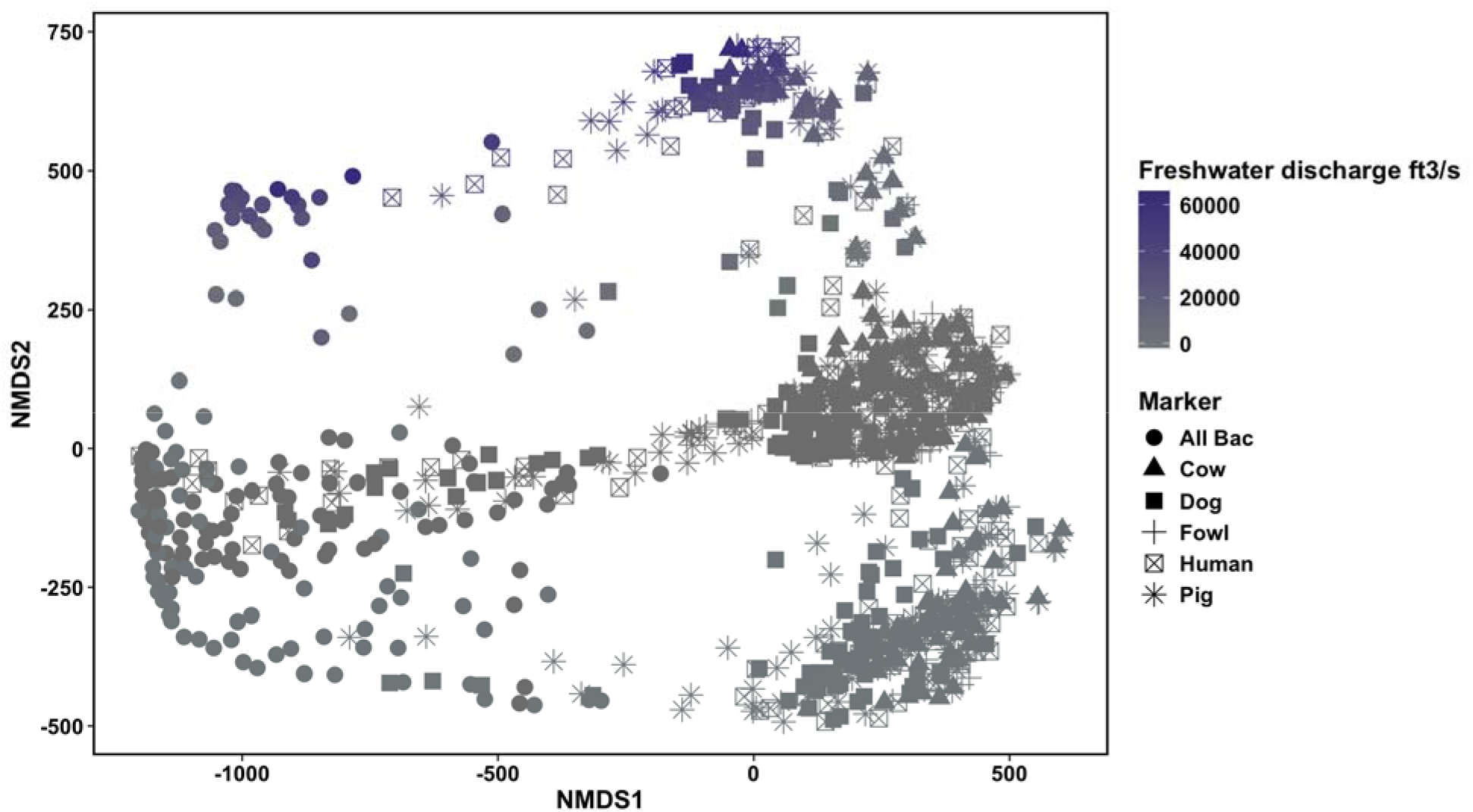
*Bacteroides* gene copy number ordinations (NMDS) constructed from freshwater discharge measurements. Points are colored by discharge and shape denotes *Bacteroides* marker.

The PCA of *Bacteroides* gene copy numbers and water chemistry parameters showed a tight cluster of all sites, with the exception of WMB, which was separated from the other sites (**Fig. 6**) due to differences in nutrient concentrations (TDN, nitrate, nitrite, ammonium).

**Figure 6.**
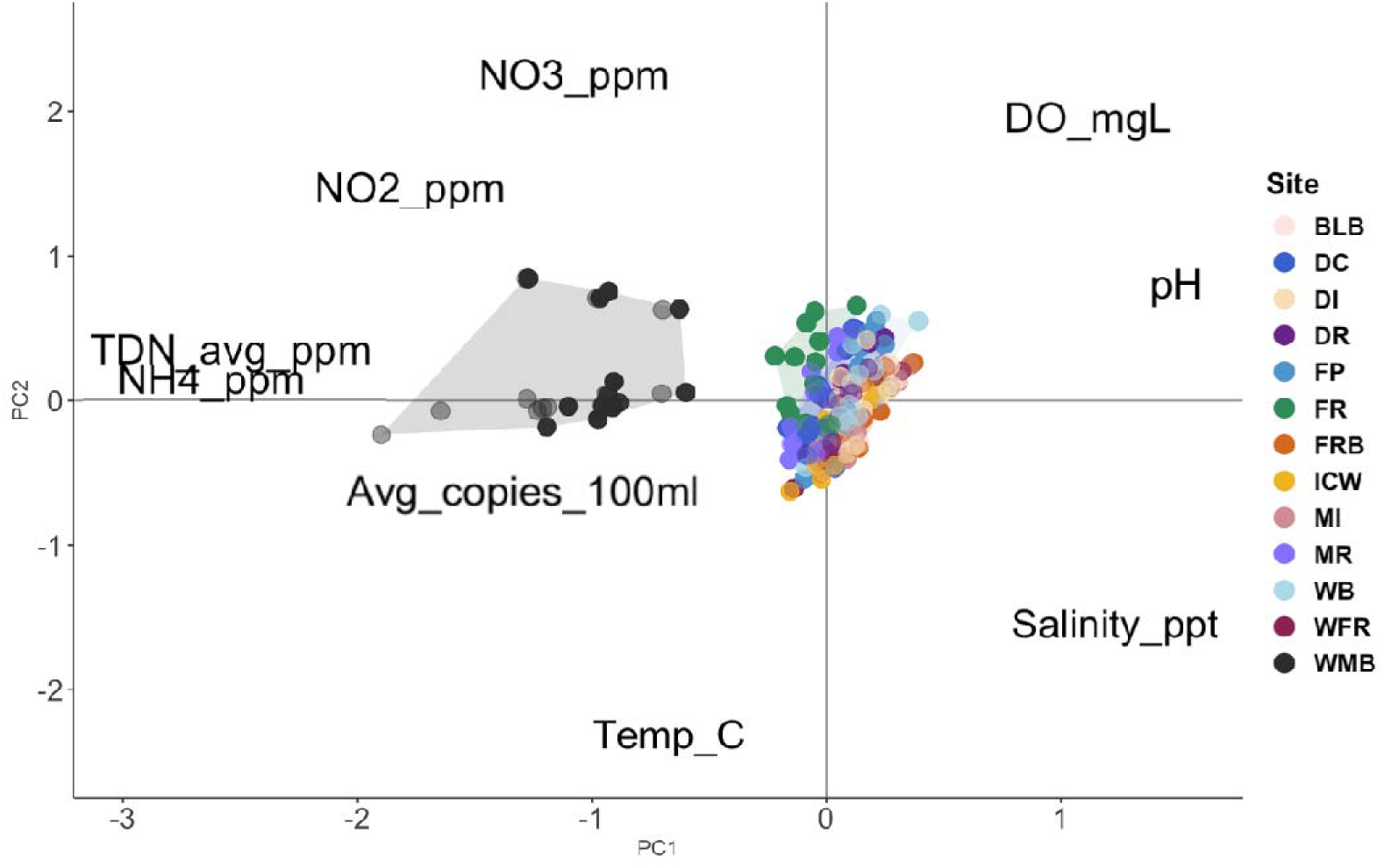
*Bacteroides* gene copy number ordinations (PCA) constructed from water chemistry and nitrogen measurements. *Bacteroides* gene copy numbers from the WMB (wastewater effluent) site clustered separately from the rest of the sites, driven by nitrogen measurements.

## Discussion

This study is the first conventional qPCR fecal source tracking effort to simultaneously validate five species from the same water sample. An additional aim of this study was to provide a baseline of fecal contamination origins and concentrations in the urbanized Mobile Bay estuary, which was accomplished across spatial and temporal scales. To do this, six qPCR primer sets targeting *Bacteroides* (AllBac, HF183/BacR287, Pig-2-Bac, DG3, Chicken/Duck-Bac, and CowM2) were verified and optimized for use within this system. The efficacy of source tracking qPCR primer sets is dependent on geographic location, and species-specific markers have displayed significant cross-reactivity (Ahmed et al. 2016). The primer sets used in this study were sensitive and specific for feces collected in the Mobile Bay watershed and did not cross-react.

Overall, human-associated *Bacteroides* were a consistent contributor to the Mobile Bay estuary, and the human marker had the strongest relationship to the total (AllBac) marker. General, human-, pig-, and dog-associated *Bacteroides* were positively related with percent impervious surface cover, and all markers had a relationship with developed land use. These findings are consistent with previous studies that found impervious surface cover is an important factor associated with higher abundances of fecal bacteria in surface waters (Mallin et al. 2000). Impervious surfaces (such as roads, sidewalks, and parking lots) reduce soil infiltration and concentrate and convey runoff to receiving waters (Miller et al. 2014). Soil infiltration decreases the transport of nonpoint source contaminants to surface water bodies, by capturing contaminants and decreasing the volume available for runoff (Weiss et al. 2008, Lambert et al. 2021). The land surrounding Mobile Bay is largely developed, specifically in the north Bay (Mobile and Daphne areas). North Mobile Bay also has wastewater infrastructure that is outdated and under supported, such as in the areas adjacent to the Mobile River (MR) sampling site. The high degree of development and large human population combined with under supported infrastructure likely accounts for the greater concentrations of fecal-associated bacteria (both general and human-associated) at these northern sites.

Dog-and pig-associated *Bacteroides* were major contributors to the system but were more geographically localized and seasonally inconsistent compared to human-associated. When averaged over all timepoints, elevated levels of dog-associated *Bacteroides* compared to other *Bacteroides* markers were observed in the Eastern Mississippi Sound sites (MI, FRB, and WFR). Urban areas often have a greater presence of canine feces, and it is a common habit for dog owners to not remove feces from streets, parks, etc. (Cinquepalmi et al. 2013, Rubel and Carbajo 2019). Dog feces left on the ground can be transported to waterways through runoff and heavy rain events, which may be exacerbated after dry periods (Li & Barrett 2008, Tian et al. 2009) and when feces are left on impervious surfaces (Mallin et al. 2000, Miller et al. 2014). The eastern sites experienced elevated dog inputs especially in the fall months (November and December), which are characterized by less intense precipitation (data not shown). The Mobile Bay estuary receives the least amount of precipitation during the month of October (both historically and during our sample collection) (NOAA), which would explain an increased flushing of canine feces after a dry period accumulation. Additionally, the occurrence and duration of dog walking can be influenced by weather conditions. For example, hot weather can reduce dog walking (Hall et al. 2021), which may be reflected in increased canine fecal concentrations seen in the cooler fall months.

Average pig-associated *Bacteroides* contributions were prominent at sites on the eastern coast of Mobile Bay (DC, FP, WB, FR, ICW), where there are few agricultural pig farms. The Pig-2-Bac marker amplifies feces from domestic pigs (Mieszkin et al. 2009) as well as wild hogs (*Sus scrofa*, this study), a species invasive to the Mobile Bay area that poses threats to the economy and public health (Fern et al. 2021). Previous studies have found wild pigs to decrease water quality, through bioturbation and fecal contributions (Kaller & Kelso 2003, Gray et al. 2019, McKee et al. 2021). Wild pigs have been demonstrated to be contributors of nonpoint fecal contamination, with variation in contributions due to their movement among watersheds and their behavioral changes in response to environmental (seasonal, resource needs) and human pressures (Bradley et al. 2025). High pig inputs were detected at the east Bay sites in May and June (data not shown), which may correspond with farrowing (birthing) peaks and greater pig populations (Sweeney et al. 1979). Besides those timepoints, there were other elevated pig marker concentrations that were localized and not seen across sites. These “spikes” in pig-associated fecal markers may represent behavioral variability in this nonpoint origin of contamination.

The AllBac marker targets the genus *Bacteroides* and has been used as a measure of “total” fecal contamination (Layton et al. 2006, Ravaliya et al. 2014, Jamison et al. 2022). In this study, the AllBac marker was detected at higher concentrations and more consistently than any other (species-specific) marker. However, the sum of species-specific marker gene copy numbers did not account for a significant portion of AllBac copy numbers for any sample (with the exception of WMB in July 2023). In a few cases, this discrepancy has been attributed to other potential fecal contributors that were not included in that particular study (Xue et al. 2018, Pendergraph et al. 2021). Our study accounted for human, dog, pig, cow, and fowl, which are major terrestrial populations in this system. We posit that this discrepancy between the AllBac marker and species-specific marker concentrations would not be accounted for by including missing major fecal contributors in the study design (e.g., felines, other avians); instead, AllBac may be amplifying *Bacteroides* species that are not explicitly associated with feces (Diston et al. 2015), such as algal-associated and free living *Bacteroides*. Additionally, *Bacteroides* species are not relegated to the guts of terrestrial organisms, they have also been found in the microbiome of aquatic organisms, such as fish (Kabiri et al. 2013). Therefore, AllBac may not be an accurate estimation or guide for determining total terrestrial fecal input to a system. Moreover, the AllBac marker targets the 16S rRNA gene and not all published *Bacteroides* qPCR assays target ribosomal gene markers, further contributing to inconsistencies between the sum of the species-specific markers and AllBac. The human, pig, and fowl markers used in this study are ribosomal, while the cow and dog markers used in this study are non-ribosomal. Lower abundance of the CowM2 marker in comparison to 16S rRNA gene markers has previously been established (Shanks et al. 2008) and may have contributed to the low detection levels of cow-associated *Bacteroides* in this study. A prior study in Fairhope, Alabama (eastern Mobile Bay) that utilized the CowM3 marker found low levels of cow-associated *Bacteroides* (they detected cow in 2% of samples), similar to this study (Xue et al. 2018). In a study comparing qPCR detection and quantification limits for the CowM2 and CowM3 markers, the CowM3 marker outperformed CowM2 (Xue and Feng 2019). However, we found that CowM3 consistently amplified wild hog feces in the current study while CowM2 did not (data not shown). Therefore, the CowM2 marker was used in this study to ensure specificity of the markers. Inconsistencies in the abundance (and subsequent amplification) of non-ribosomal and 16S rRNA gene markers may also explain why the dog marker copy number exceeded that of AllBac for WMB in July 2023. The use of AllBac as an estimate of total fecal input may need to be reevaluated, especially when using a suite of markers that include ribosomal and non-ribosomal targets.

Mobile Bay typically experiences seasonality from higher freshwater discharge in winter and spring and lower discharge in summer and fall (Stumpf et al. 1993, Du et al. 2018). The sampling period for this study occurred during a ‘low’ discharge period based on historical discharge rates for Mobile Bay (Kim and Park 2012; Du et al. 2018, Yuhl 2025). Therefore, there may not have been sufficient seasonal variation during this sampling period to capture a ‘typical’ range of seasonal differences for this system. Even under low discharge conditions, freshwater discharge still exhibited a positive relationship with *Bacteroides* gene copy numbers (AllBac and human-associated). This is consistent with previous studies in the area that have suggested that high runoff (from increased precipitation and freshwater discharge during the cold season) and high flow river systems are important contributors of feces (Gancel 2000, Frith et al. 2021). Lower salinity (YSI measurements and Sr:Ba), indicative of greater freshwater input, also exhibited relationships with higher gene copy numbers for AllBac, human, fowl, and cow markers. Further research is necessary to better understand the effects of higher freshwater discharge periods and more prominent seasonality in the Mobile Bay estuary.

All of the *Bacteroides* markers showed negative relationships with δ^15^N and positive relationships with total dissolved nitrogen (TDN) and at least one other nitrogen species. Statistical model selection attributed TDN as the best predictor of *Bacteroides* copy numbers. Lower δ^15^N values are associated with raw sewage and high-flow discharge from wastewater treatment plants in populated areas, including Mobile Bay (Daskin et al. 2008, Biancani et al. 2012), whereas greater δ^15^N is associated with processed nitrogen in treated sewage (McClelland et al. 1997, Savage 2005, Loomer et al. 2015, Darrow et al. 2016). The combination of qPCR markers and stable isotope indicators revealed that increased fecal inputs to this system are likely from raw sewage and high-flow treated wastewater discharge, rather than groundwater infiltrated or processed waste that includes secondary nitrogen removal steps (Archana et al. 2016, Wang et al. 2024). Raw sewage, in particular, can be linked to under supported municipal (e.g., MAWSS) and private (septic, dumping from boats and homes) infrastructure, combined sewer overflows (CSO), and various terrestrial sources. The Mobile Bay area has a combination of sewer and septic systems, especially in the more developed northern areas, and spills are not uncommon (ADEM Sanitary Sewer Overflow eFile portal). Heavy rains (characteristic of the Mobile Bay system) can cause CSO (Mailhot et al. 2015, MAWSS 2025) when treatment capacity is exceeded and aging sewage infrastructure is prone to failure and accidents (Fenner 2000, Brown & Peterson 2012). Additionally, ongoing construction can remobilize fecal indicators (Frith et al. 2021). Hence, a combination of aging infrastructure, sewage spills, and leaks likely contribute to elevated levels of human-associated fecal matter in Alabama’s coastal waters.

Samples collected at the effluent pipe of the major wastewater treatment plant serving much of the eastern shore of Mobile Bay (WMB) had consistently greater concentrations of *Bacteroides* and nitrogen constituents than other sites. This finding is not surprising because these samples were collected before the treated effluent mixed with the receiving waterbody. Other samples representative of wastewater effluent (BLB, DC) were collected near outfall pipes but after the effluent entered and mixed with the receiving waters (diluted effluent compared to WMB). This sampling difference may explain the distinction found between the WMB effluent *Bacteroides* and these other sites, driven by higher concentrations of nitrogen constituents (**Fig. 6**). Prior studies have also found high NH_4_ ^+^ and low δ^15^N at this location (Daskin et al. 2008, Biancani et al. 2012, Gancel 2020), as well as higher concentrations of indicator organisms compared to low flow wastewater treatment systems in the area (BLB) (Gancel 2020). The relationship between elevated *Bacteroides* and nitrogen concentrations (especially TDN and NH_4_ ^+^) in the WMB samples can serve as a guide for identifying human sewage signatures in coastal waterways. TDN and NH_4_ ^+^ can potentially serve as reliable predictors of human fecal inputs for future studies.

This study sets a baseline for temporal and spatial variation in biological sources of fecal contamination to an urbanized coastal estuary. Human-associated feces were a consistent major contributor to the system, with localized inputs from pig (wild or domestic) and dog, indicating potential for wildlife and domestic animal contributions that may vary locally depending on land use. Freshwater discharge and nitrogen constituents were good predictors of *Bacteroides* concentrations in this system, while areas with impervious surfaces and developed land cover likely contributed to elevated fecal indicator levels by facilitating conveyance from the watershed. These findings highlight the need to support regular monitoring, maintenance, repair, and proper capacity of wastewater treatment facilities to meet the needs of growing communities, particularly in coastal areas that face increasing pressures from changing climate and storm intensity. Importantly, this study demonstrates successful verification and use of multi-species molecular MST methods that can be readily applied in other watersheds, leading to the development of better detection, identification, and management strategies for community health of coastal areas globally.

## Acknowledgements

We appreciate our stakeholders for their continuous support. We thank Kristin DeRosia-Banick for her GIS skills and the Carmichael lab members who helped us during our boat sampling. We thank Dr. Jeremiah Henning and Dr. Amy Willis (STAMPS) for their statistics knowledge and guidance. This work was funded by Environmental Protection Agency grant MX-02D17922 to BKR, RHC, and SNC and the Dauphin Island Sea Lab/ U.S. Food and Drug Administration Joint Graduate Research Program (5U19FD005923-09).

