## Supplemental for "Fecal Contributors to an Urbanized Estuary Identified by a Multi-Species qPCR Approach"

**Appendixes**

Table A1. Optimized master mix composition for qPCR primer sets. ROX (passive reference dye) was added at a volume of 0.06μl per reaction for all assays and 2μl of target DNA template was used in each reaction. The total reaction volume was brought to 25μl with nuclease-free water. Invitrogen Platinum Taq DNA polymerase was used for all reactions (Thermo Fisher Scientific, Waltham, Massachusetts). *Bacteroides* primers and probes are denoted by Bac and internal amplification control primers and probe are denoted by IAC.

| **Primer Set** | **[Final]** | | | | | | | | | |
| --- | --- | --- | --- | --- | --- | --- | --- | --- | --- | --- |
|  | **Buffer** | **MgCl_2_** | **dNTPs** | **Forward primer (Bac)** | **Reverse primer (Bac)** | **Forward primer (IAC)** | **Reverse primer (IAC)** | **Probe (IAC)** | **Probe (Bac)** | **Taq** |
| AllBac | 1.0 | 5.0 | 0.4 | 0.2 | 0.2 | 0.1 | 0.1 | 0.15 | 0.1 | 2.0 |
| Human | 1.0 | 3.0 | 0.4 | 1.0 | 1.0 | 0.2 | 0.2 | 0.15 | 0.08 | 2.0 |
| Pig | 1.0 | 5.0 | 0.4 | 0.2 | 0.2 | 0.2 | 0.2 | 0.15 | 0.1 | 2.0 |
| Cow | 1.0 | 3.0 | 0.4 | 1.0 | 1.0 | 0.2 | 0.2 | 0.15 | 0.08 | 2.0 |
| Dog | 1.0 | 3.0 | 0.4 | 1.4 | 1.4 | 0.2 | 0.2 | 0.15 | 0.1 | 2.0 |
| Fowl | 1.0 | 3.0 | 0.4 | 0.2 | 0.2 | 0.2 | 0.2 | 0.15 | 0.2 | 2.0 |

Table A2. Optimized cycling conditions for qPCR assays. All assays had an initial denaturation step of 95^o^C for 120 seconds and were run for 40 cycles.

| **Primer Set** | **Denaturation** | | **Anneal/Extend** | |
| --- | --- | --- | --- | --- |
|  | **Temp (^o^C)** | **Time (s)** | **Temp (^o^C)** | **Time (s)** |
| AllBac | 95 | 15 | 60 | 60 |
| Human | 95 | 15 | 60 | 60 |
| Pig | 95 | 15 | 60 | 60 |
| Cow | 95 | 15 | 60 | 60 |
| Dog | 95 | 15 | 60 | 60 |
| Fowl | 95 | 5 | 60 | 31 |

Table A3. Average and standard error of environmental parameter measurements at each site over all timepoints. Water temperature, salinity, dissolved oxygen, and pH were collected a handheld YSI during sampling. Discharge data compiled from USGS databases, turbidity data compiled from ARCOS, and precipitation data compiled from South Alabama Mesonet.

| Site | Precipitation (24 hrs) | Water Temp (°C) | Salinity | DO (mg/L) | pH | NO_3_^-^ (ppm) | NO_2_^-^ (ppm) | NH_4_^+^ (ppm) | TDN (ppm) | δ^15^N_Air_ | Sr:Ba | Two week avg discharge (ft^3^/s) | Two week avg turbidity (NTU) |
| --- | --- | --- | --- | --- | --- | --- | --- | --- | --- | --- | --- | --- | --- |
| BLB | 0.002±0.001 | 24.2±1.6 | 17.5±1.6 | 7.4±0.3 | 7.5±0.1 | 0.2±0.1 | 0.006±0.005 | 0.28±0.08 | 0.34±0.06 | 6.7±0.4 | 41.1±5.9 | - | - |
| MI | 0.002±0.001 | 25.2±1.8 | 15.9±2.1 | 7.4±0.4 | 7.7±0.1 | 0.02±0.01 | 0.002±0.002 | 0.13±0.02 | 0.20±0.02 | 6.5±0.1 | 44.0±7.2 | - | - |
| FRB | 0.002±0.001 | 24.4±1.7 | 15.3±1.9 | 7.1±0.4 | 7.7±0.1 | 0.02±0.01 | 0.0003±0.0002 | 0.1±0.01 | 0.2±0.01 | 6.1±0.2 | 36.3±5.9 | 43.8±8.5 | 26.8±5.3 |
| WFR | 0.002±0.001 | 24.5±1.7 | 11.7±1.9 | 6.1±0.4 | 7.4±0.1 | 0.07±0.03 | 0.002±0.0007 | 0.1±0.01 | 0.27±0.02 | 5.4±0.2 | 25.7±4.1 | 43.8±8.5 | 26.8±5.3 |
| DR | 0.002±0.001 | 24.1±1.8 | 5.0±1.0 | 7.5±0.4 | 7.6±0.1 | 0.09±0.04 | 0.003±0.001 | 0.05±0.009 | 0.23±0.01 | 7.1±0.2 | 13.1±2.3 | - | 55.7±25.2 |
| WMB | 0.002±0.001 | 25.9±1.0 | 0.5±0.04 | 5.8±0.2 | 7.0±0.1 | 1.3±0.4 | 1.1±0.4 | 15.6±1.6 | 11.1±0.8 | 1.6±0.3 | 1.4±0.2 | - | - |
| MR | 0.003±0.002 | 23.4±1.8 | 6.5±1.6 | 6.6±0.4 | 7.1±0.1 | 0.9±0.1 | 0.03±0.007 | 0.1±0.01 | 0.4±0.02 | 4.6±0.6 | 17.0±4.6 | 30097±4864 | 55.7±25.2 |
| DC | 0.002±0.001 | 23.2±1.5 | 0.5±0.4 | 6.1±0.4 | 7.7±0.1 | 0.8±0.09 | 0.007±0.0007 | 0.08±0.008 | 0.3±0.02 | 3.2±0.4 | 1.3±0.8 | - | 9.0±1.3 |
| FP | 0.002±0.001 | 23.8±1.9 | 8.1±1.7 | 7.3±0.5 | 7.6±0.1 | 0.2±0.08 | 0.005±0.002 | 0.06±0.01 | 0.2±0.02 | 7.5±0.3 | 16.3±3.2 | 26983±4791 | - |
| WB | 0.001±0.0008 | 24.0±1.8 | 8.9±1.2 | 8.2±0.4 | 7.7±0.1 | 0.3±0.1 | 0.005±0.002 | 0.07±0.009 | 0.3±0.02 | 6.9±0.1 | 18.6±2.6 | 193.8±22.7 | 17.8±1.9 |
| FR | 0.001±0.0008 | 24.2±1.5 | 4.9±1.0 | 7.2±0.5 | 7.4±0.1 | 2.1±0.3 | 0.02±0.003 | 0.06±0.009 | 0.6±0.05 | 6.4±0.1 | 8.7±2.0 | 355.8±44.5 | 33.1±11.5 |
| ICW | 0.001±0.0008 | 24.5±1.6 | 14.5±1.3 | 5.6±0.4 | 7.2±0.1 | 0.07±0.02 | 0.003±0.0009 | 0.1±0.01 | 0.2±0.02 | 6.6±0.1 | 29.5±3.6 | 556±130 | - |
| DI | 0.002±0.001 | 22.9±1.7 | 15.6±2.2 | 7.2±0.4 | 7.5±0.1 | 0.2±0.06 | 0.008±0.002 | 0.1±0.01 | 0.2±0.02 | 6.6±0.1 | 35.6±6.1 | - | 21.4±5.3 |

Table A4. Results of dredge and model.sel functions to determine the best predictor of *Bacteroides* copy numbers. Environmental parameters with missing values were omitted from the analysis. Models ranked by AICc(x).

|  | Salinity | TDN | NH_4_^+^ | NO_2_^-^ | NO_3_^-^ | DO | Land Use |
| --- | --- | --- | --- | --- | --- | --- | --- |
| Sum of weights (dredge) | 0.97 | 0.58 | 0.60 | 0.35 | 0.30 | 0.29 | 0.03 |
| Weight (model.sel) | 0.0 | 0.988 | 0.012 | 0.0 | 0.0 | 0.0 | 0.0 |

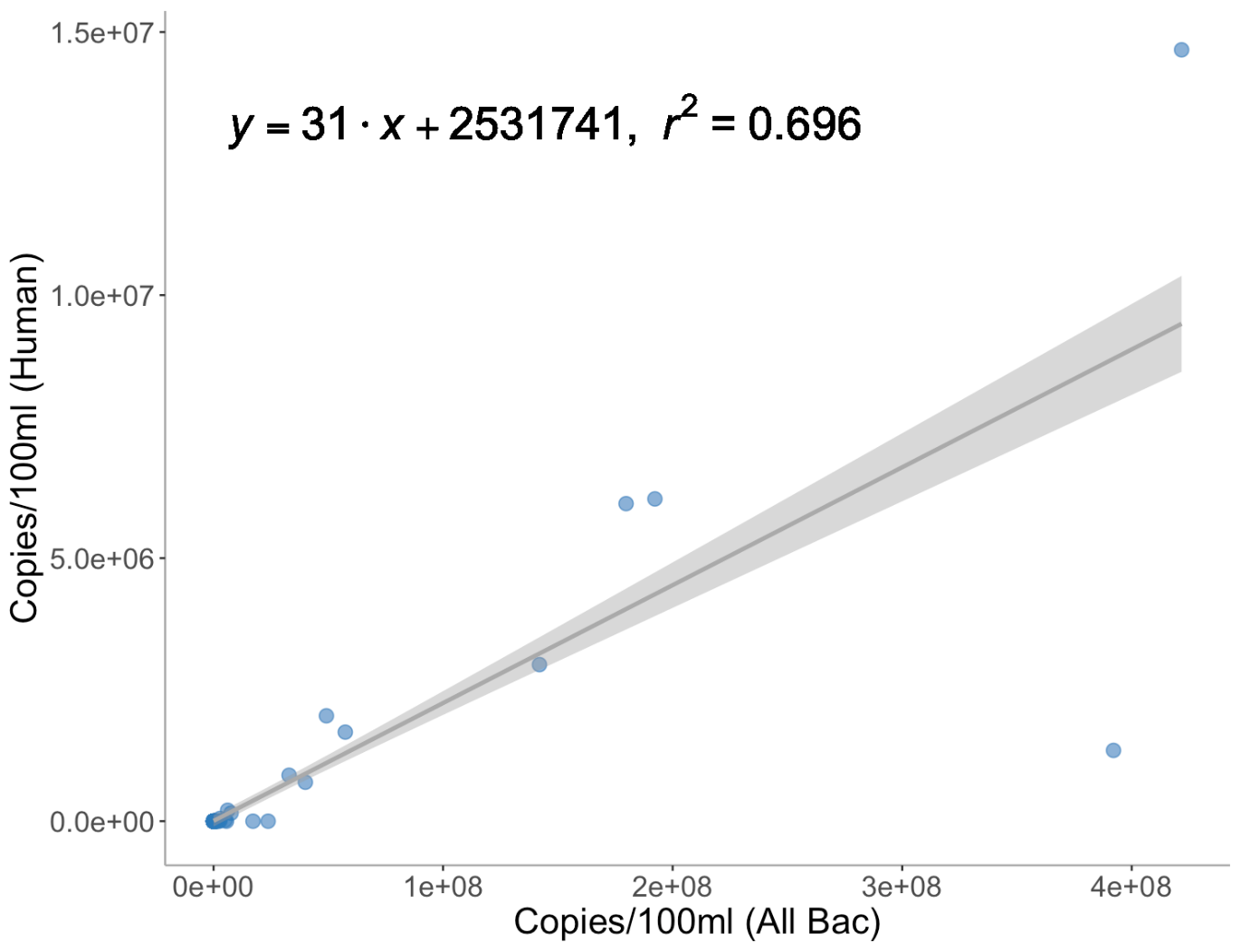

Figure A1. Significant positive correlation between human-specific and general *Bacteroidales* (AllBac) gene markers (r2=0.696, p<2.2e-16).

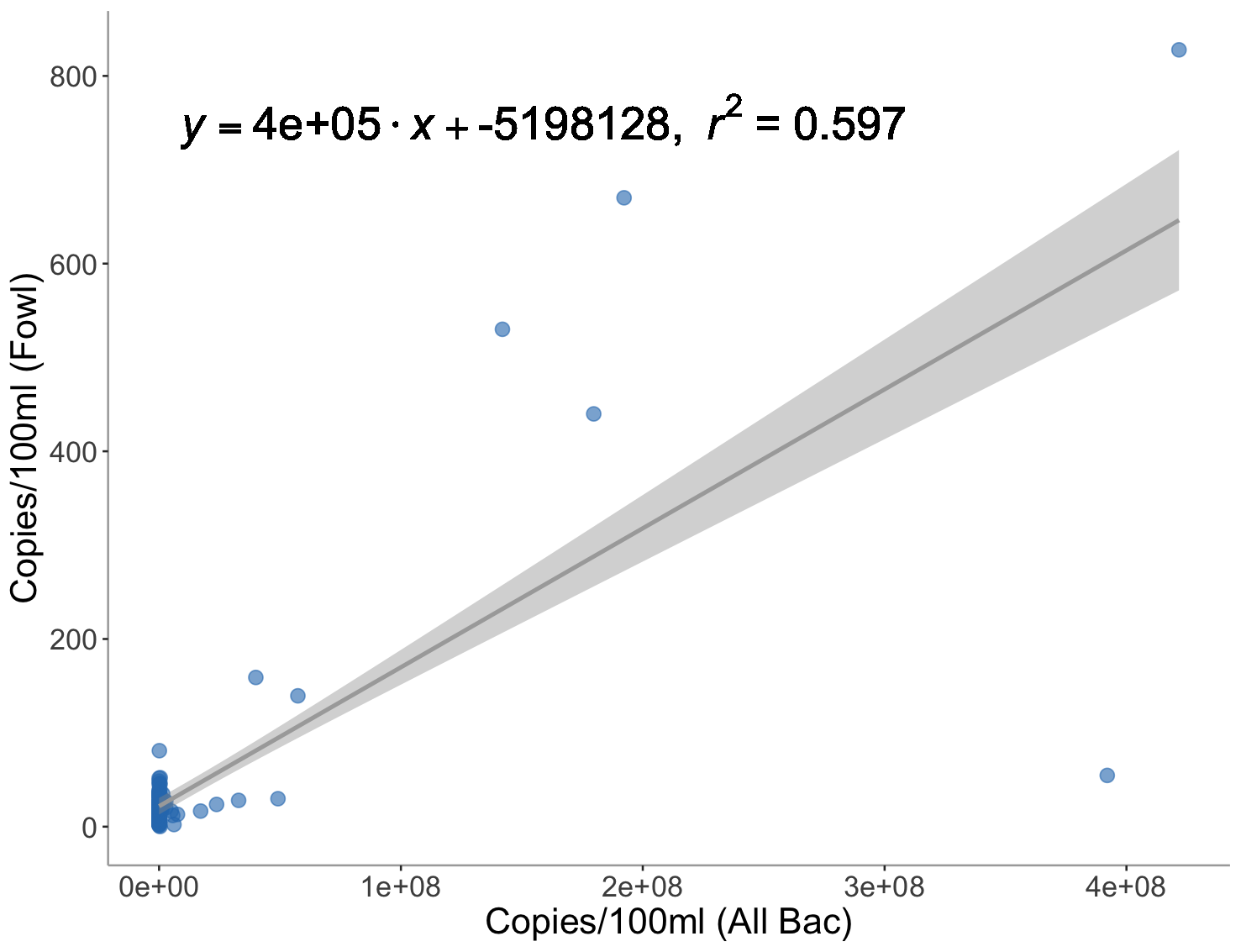

Figure A2. Significant positive correlation between fowl-specific and general *Bacteroidales* (AllBac) gene markers (r2=0.597, p<2.2e-16).

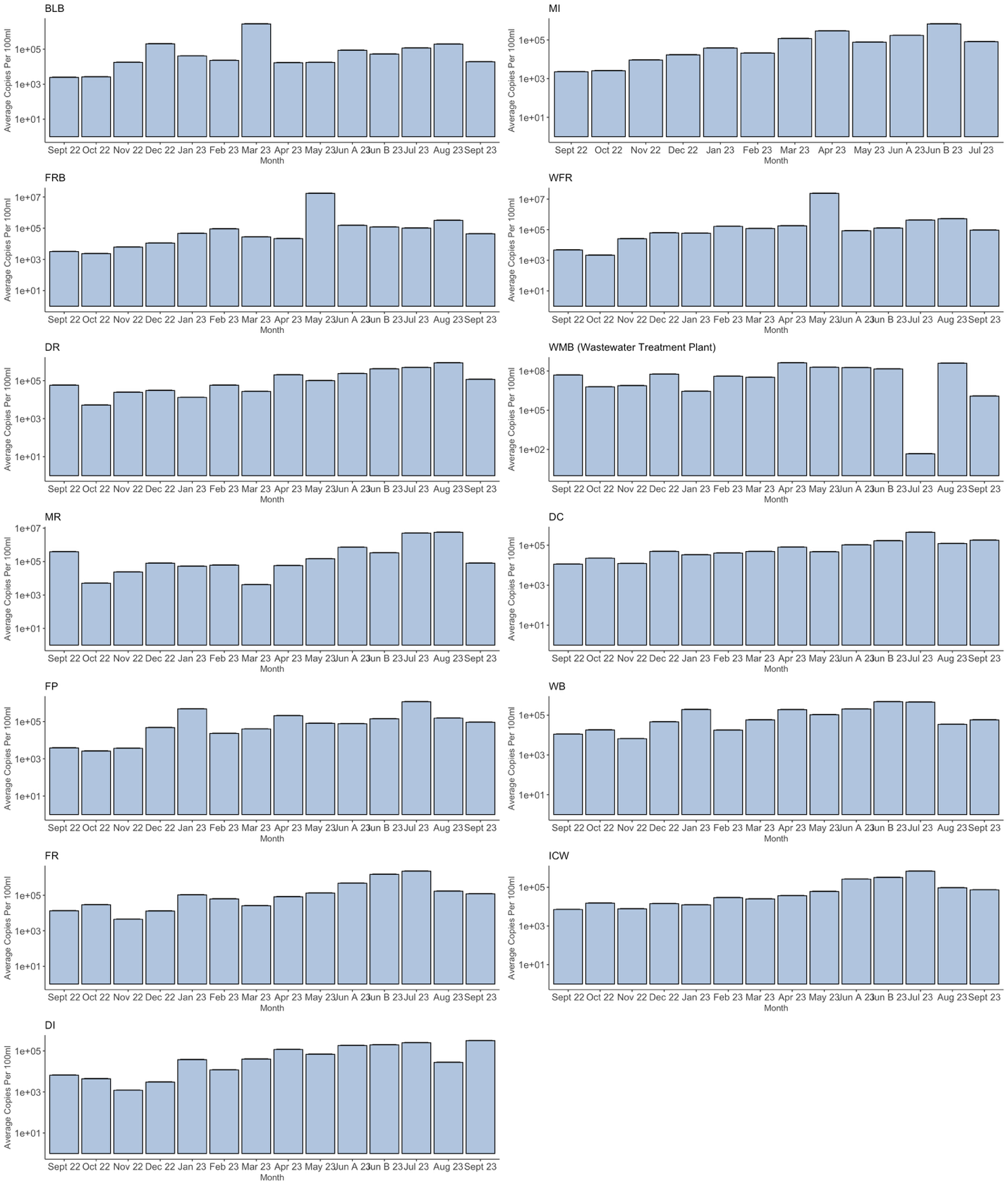

Figure A3. AllBac gene copy numbers (per 100ml) at each site over all timepoints. Axis scales are not consistent between sites.
